# Airway Gene-Expression Classifiers for Respiratory Syncytial Virus (RSV) Disease Severity in Infants

**DOI:** 10.1101/628701

**Authors:** Lu Wang, Chin-Yi Chu, Matthew N. McCall, Christopher Slaunwhite, Jeanne Holden-Wiltse, Anthony Corbett, Ann R. Falsey, David J. Topham, Mary T. Caserta, Thomas J Mariani, Edward E. Walsh, Xing Qiu

**Author notes:** These authors contributed equally to this work (Lu Wang, Chin-Yi Chu). Corresponding authors: Thomas J. Mariani; Edward E. Walsh; Xing Qiu.

## Abstract

**Background:** A substantial number of infants infected with RSV develop severe symptoms requiring hospitalization. We currently lack accurate biomarkers that are associated with severe illness.

**Method:** We defined airway gene expression profiles based on RNA sequencing from nasal brush samples from 106 full-tem previously healthy RSV infected subjects during acute infection (day 1-10 of illness) and convalescence stage (day 28 of illness). All subjects were assigned a clinical illness severity score (GRSS). Using AIC-based model selection, we built a sparse linear correlate of GRSS based on 41 genes (NGSS1). We also built an alternate model based upon 13 genes associated with severe infection acutely but displaying stable expression over time (NGSS2).

**Results:** NGSS1 is strongly correlated with the disease severity, demonstrating a naïve correlation (ρ) of ρ=0.935 and cross-validated correlation of 0.813. As a binary classifier (mild versus severe), NGSS1 correctly classifies disease severity in 89.6% of the subjects following cross-validation. NGSS2 has slightly less, but comparable, accuracy with a cross-validated correlation of 0.741 and classification accuracy of 84.0%.

**Conclusion:** Airway gene expression patterns, obtained following a minimally-invasive procedure, have potential utility for development of clinically useful biomarkers that correlate with disease severity in primary RSV infection.

## Introduction

Respiratory Syncytial Virus (RSV) is the most important cause of respiratory illness in infants and young children, accounting for more than 57,000 bronchiolitis and pneumonia hospitalizations in the US annually.[1] Worldwide, 33.1 million acute lower respiratory infections and 3.2 million hospitalizations in children under 5 years of age are attributed to RSV each year.[2] In the US ~1-2% of newborns are hospitalized during their first winter, with rates greatest in the first two months of life (25.9 per 1000).[3] Risk factors for severe disease include gestational age < 29 weeks, bronchopulmonary disease and symptomatic congenital cardiac disease, while less well defined risks include lack of breast feeding, and exposure to tobacco smoke. However, the majority of hospitalized infants are full-term infants whose only risk factor is young age at the time of infection.[3]

A number of severity scores using clinical parameters, including cutaneous oximetry, have been used to grade illness severity for use in management and as an outcome in therapeutic, or potentially, vaccine trials. [4–13] However, none of the clinically based severity scores have been universally adopted.[14] Reasons may include heterogeneity in the scope and purpose of the score, the ages to which it is applied and concerns about inter-observer variability and subjectivity in interpreting clinical signs, including oximetry, that often are temporally dynamic over short intervals. Identification of an objective biomarker that accurately correlates with, or potentially predicts, disease severity could be highly useful.[15, 16]

We and others have reported a relationship between disease severity and host gene expression in peripheral blood cells and nasal swab samples during infection.[17–20] These results suggest such an approach may allow development of biomarkers to accurately categorize RSV disease severity. As part of the AsPIRES study[21] we recently reported on the feasibility of measuring gene expression of airway cells collected by nasal swab in healthy infants in order to study RSV disease pathogenesis.[22] However, in this manuscript, we describe the use of this gene expression data during RSV infection to develop two airway gene expression-based classifiers that are highly correlated with clinical disease severity. This represents a first step in developing a biomarker using gene expression responses capable of accurately classifying clinical severity in primary RSV-infection that could be used in conjunction with clinical evaluation.

## Methods

### Study Subjects

Subjects included RSV infected infants enrolled in the AsPIRES study at the University of Rochester Medical Center (URMC) and Rochester General Hospital (RGH).[21] The Institutional Review Boards of both institutions approved the study, and all parents provided written informed consent. RSV-infected infants came from three cohorts during three winters (October 2012 through April 2015); one cohort included infants hospitalized with RSV, a second cohort was recruited at birth and followed through their first winter for development of RSV infection, and the third cohort was RSV infected infants seen in pediatric offices and emergency departments and managed as outpatients. All subjects were full-term infants undergoing a primary RSV infection during their first winter season. Nasal samples were collected from the inferior nasal turbinate, by gentle brushing with a flocked swab as previously described [22], during the acute illness visit (visit 1) and at a convalescent visit ~28 after illness onset (visit 2). Illness severity was graded from 0-10 using a Global Respiratory Severity Score (GRSS), that uses nine parameters (age adjusted respiratory rate, chest retractions, wheezing, rales/rhonchi, apnea, cyanosis, room air oxygen saturation, lethargy and poor feeding) as previously described.[23] We defined a GRSS >3.5 as severe disease as it is highly correlated with illness requiring hospitalization.

### Nasal RNA processing

The process for nasal RNA recovery was previously described.[22] Briefly, following flushing of the nares with 5 milliliters of saline to remove mucus and cellular debris, a flocked swab was used to recover cells at the level of the turbinates. The swab was immediately placed in RNA stabilizer (RNAprotect, Qiagen, Germantown, MD) and maintained at 4 °C. Cells were recovered by filtering through a 0.45 uM membrane filter. Recovered cells were lysed and homogenized using the AbsolutelyRNA Miniprep kit (Agilent, Santa Clara, CA) according to the manufacturer’s instructions. 1 ng of total RNA was amplified using the SMARter Ultra Low amplification kit (Clontech, Mountain View, CA) and libraries were constructed using the NexteraXT library kit (Illumina, San Diego, CA). Libraries were sequenced on the Illumina HiSeq2500. Sequences were aligned against human genome version of hg19 using STARv2.5, counted with HTSeq, and normalized by Fragments Per Kilobase of transcript per Million mapped reads (FPKM). A total of 6,844 transcription profiles (genes) were reported after quality assurance analysis and preprocessing. Additional technical details on data preprocessing can be found in Supplementary Text.

### Statistical methods

Descriptive statistics are reported in Table 1. Discrete variables are summarized in percentages, and continuous variables were summarized as Mean (SE). For continuous variables, we performed 2-sample Welch t-tests to check the equality between the mild and severe groups; for categorical variables, Fisher’s exact test was used instead. The nasal gene-expression severity scores we developed in this study were based on multivariate regression analysis with bi-directional stepwise model selection based on Akaike Information Criterion (AIC). Technical details of model development and cross-validation (CV) can be found in Supplementary Material. All analyses were conducted using SAS 9.3 (SAS Institute Inc., Cary, NC, USA) and the R programming language (version 3.5, R Foundation for Statistical Computing, Vienna, Austria).

**Table 1.** Demographic data of subjects. *P*-values reported in the last column were either based on Fisher’s exact test (if the variable is categorical) or Welch *t*-test (if the variable is continuous).

|  | mild (n=42)<sup>a</sup> |  | severe (n=64)<sup>a</sup> |  | p value |
| --- | --- | --- | --- | --- | --- |
|  | n | Mean (SE)<br>or % | n | Mean (SE)<br>or % |  |
| Global Severity Score | 42 | 1.63(0.15) | 64 | 6.13(0.22) | <0.001 |
| Visit Age (months) | 42 | 3.52(0.31) | 64 | 3.24(0.3) | 0.5122 |
| Gestational Age (weeks) | 42 | 39.05(0.19) | 64 | 38.8(0.18) | 0.3437 |
| Birth Weight (kg) | 42 | 3.32(0.11) | 64 | 3.36(0.07) | 0.7468 |
| Family Size | 42 | 4.43(0.44) | 64 | 3.98(0.22) | 0.3703 |
| Days Since Disease Onset | 42 | 4.31(0.27) | 64 | 4.86(0.22) | 0.1209 |
| Breast Feeding Summary | 42 | 1.56(0.19) | 63 | 1.53(0.16) | 0.8979 |
| Sex |  |  |  |  | 0.4275 |
| Male | 23 | 44.23 | 29 | 55.77 |  |
| Female | 19 | 35.19 | 35 | 64.81 |  |
| Ethnicity |  |  |  |  | 0.8018 |
| Hispanic or Latino | 8 | 42.11 | 11 | 57.89 |  |
| Non-Hispanic or Non-Latino | 34 | 39.08 | 53 | 60.92 |  |
| Race |  |  |  |  | 0.3115 |
| Caucasian | 23 | 37.1 | 39 | 62.9 |  |
| Other race | 19 | 47.5 | 21 | 52.5 |  |
| Missing | - | 0 | 4 | 100 |  |
| Delivery Type |  |  |  |  | 0.3634 |
| Vaginal | 29 | 36.71 | 50 | 63.29 |  |
| C-section | 13 | 48.15 | 14 | 51.85 |  |
| Smoking Exposure |  |  |  |  | 1 |
| Yes | 14 | 38.89 | 22 | 61.11 |  |
| No | 28 | 40 | 42 | 60 |  |
| RSV Group |  |  |  |  | 1 |
| A | 23 | 38.98 | 36 | 61.02 |  |
| B | 18 | 39.13 | 28 | 60.87 |  |
| Missing | 1 | 100 | - | 0 |  |

<sup>a</sup> based on GRSS $\leq 3.5$ (mild) or $> 3.5$ (severe)

## Results

Of the 139 RSV-infected infants enrolled in the AsPIRES study, nasal samples were available from 119 subjects during acute infection (day 1-10 of illness) and 81 subjects during convalescence (day 28 of illness). Among these 200 samples, 175 samples (106 acute samples and 69 convalescent samples) met sufficient quality to be used for subsequent analyses. Demographic and clinical information for these 106 subjects are provided in Table 1. The clinical severity score (GRSS) for these subjects ranged from 0 to 10, with 42 subjects considered to have mild disease (GRSS ≤3.5; mean ±SE GRSS of 1.63 ±0.15) and 64 to have severe disease (GRSS >3.5; mean GRSS of 6.13 ±0.22). There were no significant differences between the mild and severe groups in gender, race, delivery type, breast feeding, or exposure to tobacco smoke. There also was no difference in age at time of infection or in duration of illness at the time of evaluation.

### Nasal gene expression correlates of clinical severity during acute illness

The 6,844 genes remaining after data preprocessing and filtering were subjected to the Pearson correlation test to select genes that were significantly correlated with GRSS during acute infection. After controlling the false discovery rate (FDR) at the 0.05 level, 66 significant genes were identified.[24] Using these genes, we applied model selection procedures (see Supplementary Text for more details) to select an initial multivariate regression model for GRSS (Model 1), which was comprised of 39 genes and had relatively good predictive power (77.4% accuracy, or 24 misclassifications) for the dichotomous clinical outcome (mild vs. severe illness) in leave-one-out cross-validation (LOOCV).

Not unexpectedly, there is a strong correlation among the 66 genes, which might reduce the diagnostic performance of Model 1. Using a novel method based on principal component analysis (PCA), we identify ten supplementary genes as additional features to model GRSS (see Supplementary Text for more details). With these additional features and using the same model selection strategy, we developed two additional models: Model 2 comprised of 41 genes and Model 3 comprised of 42 genes. The performance of these models was evaluated by LOOCV (Table 2). We found that the incorporation of the supplementary genes into Model 2 (CV prediction accuracy of 89.6%; 11 misclassifications) significantly improved the accuracy compared to Model 1 (24 misclassifications) and Model 3 (23 misclassifications). Of note, Model 2 contained 5 supplementary genes, and we defined it as **NGSS1** (nasal gene expression severity score 1). As shown in Figure 1, NGSS1 is highly associative with GRSS (naïve ρ=0.935; CV ρ=0.813). For the population of subjects in the AsPIRES study, the sensitivity and specificity for identifying severe disease were high (sensitivity 90.1%, specificity 88%) which would translate to a positive predictive value (PPV) of 92% and a negative predictive value (NPV) of 86%.

**Table 2.** Performance of four models used in developing NGSS1 and NGSS2. Naïve and CV RSS are the mean residual sums of squares of the predictive model in the original and cross-validation analyses, respectively. Correlation are the Pearson correlation coefficient between the predicted severity scores and the clinically defined GRSS. Prediction accuracy is the percentage of correctly predicted mild (NGSS ≤3.5) or severe (NGSS >3.5) symptoms, compared with the same phenotype defined by the GRSS (mild: GRSS≤3.5; severe: GRSS>3.5).

|  | number of genes selected | Naïve RSS | Naïve Correlation | Naïve misclassified subjects (out of 106) | CV RSS | CV Correlation | CV prediction accuracy | CV misclassified subjects (out of 106) |
| --- | --- | --- | --- | --- | --- | --- | --- | --- |
| Model 1 | 39 genes | 1.234 | 0.909 | 15 | 2.743 | 0.797 | 77.4% | 24 |
| **Model 2\*** | **41 genes** | **0.884** | **0.935** | **9** | **2.681** | **0.813** | **89.6%** | **11** |
| Model 3 | 42 genes | 0.920 | 0.933 | 13 | 2.119 | 0.844 | 78.3% | 23 |
| **Model 4\*\*** | **13 genes** | **2.549** | **0.800** | **16** | **3.215** | **0.741** | **84.0%** | **17** |

\* designated NGSS1. \*\* designated NGSS2

**Figure 1.**
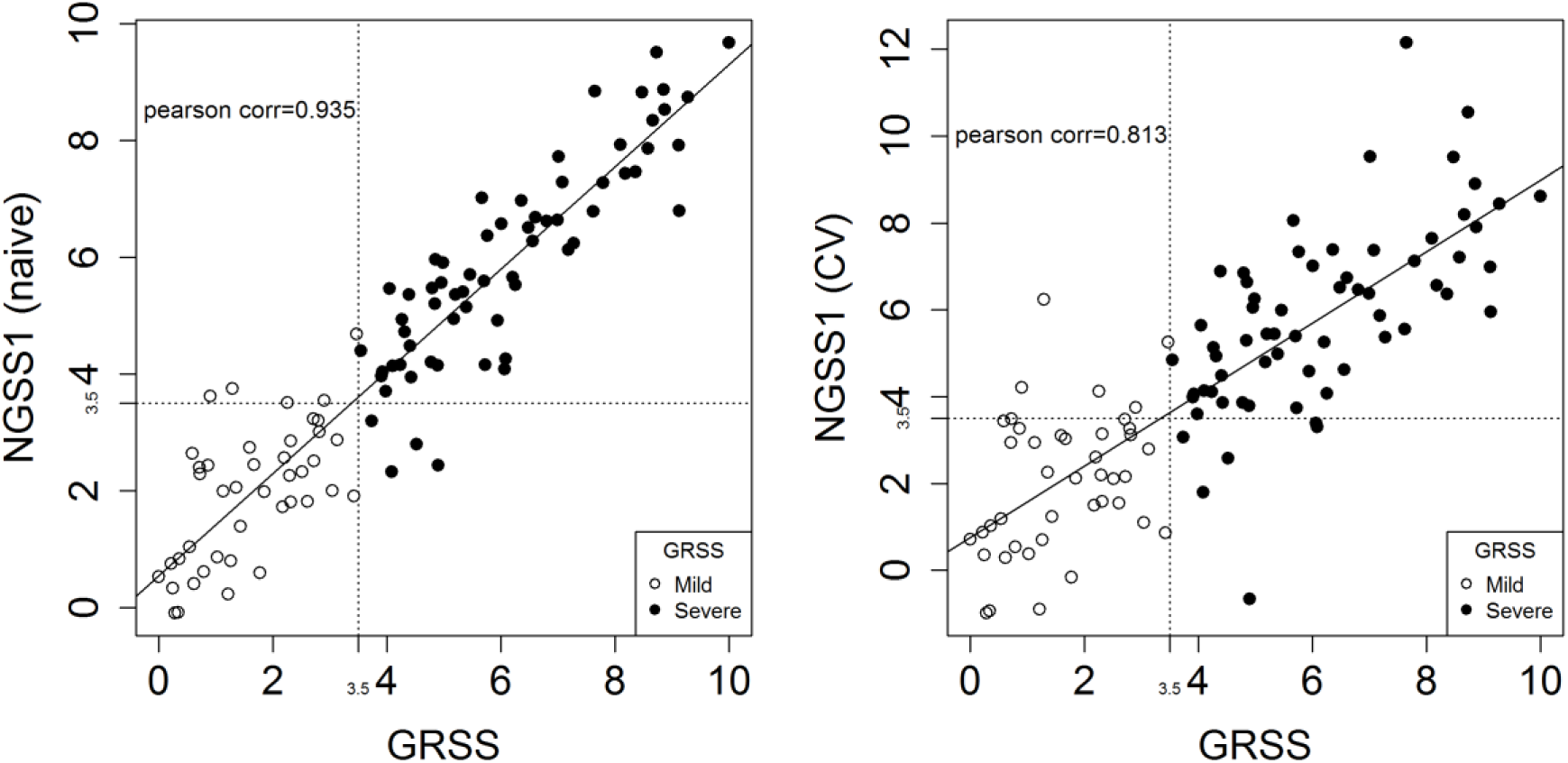
Correlating NGSS1 (severity score predicted by Model 2) with GRSS. Left: naïve Pearson correlation between GRSS and NGSS1 is ***ρ*** = **0.935**. Right: cross-validated Pearson correlation between GRSS and NGSS is ***ρ*** = **0.813**. Solid dots are subjects with severe symptoms (defined by GRSS>3.5) and empty dots are those with mild symptoms (GRSS≤3.5).

### Validation of NGSS1 at the Convalescence Phase

NGSS1 was trained exclusively from data collected at the acute phase (visit 1). For a subset (n=54) of subjects, we also had their nasal transcriptome profiles at the convalescence phase (day 28 after illness onset), a time when most infants had completely recovered from their illness. If NGSS1 is a valid surrogate for disease severity, we hypothesized that NGSS1 calculated from the severely ill subjects at visit 2 would converge to those of the mildly ill subjects. Compared with the acute visit, the calculated NGSS1 at the convalescent visit predicted a significantly lower mean severity score for severe subjects (n=29, 6.22 vs. 2.82, p<.001). In contrast, there was no significa vs. 2.31, p=0.45), nor between the severe and mild groups at visit 2 (2.82 vs. 2.31, p=0.40). These results are illustrated in Figure 2A.

**Figure 2.**
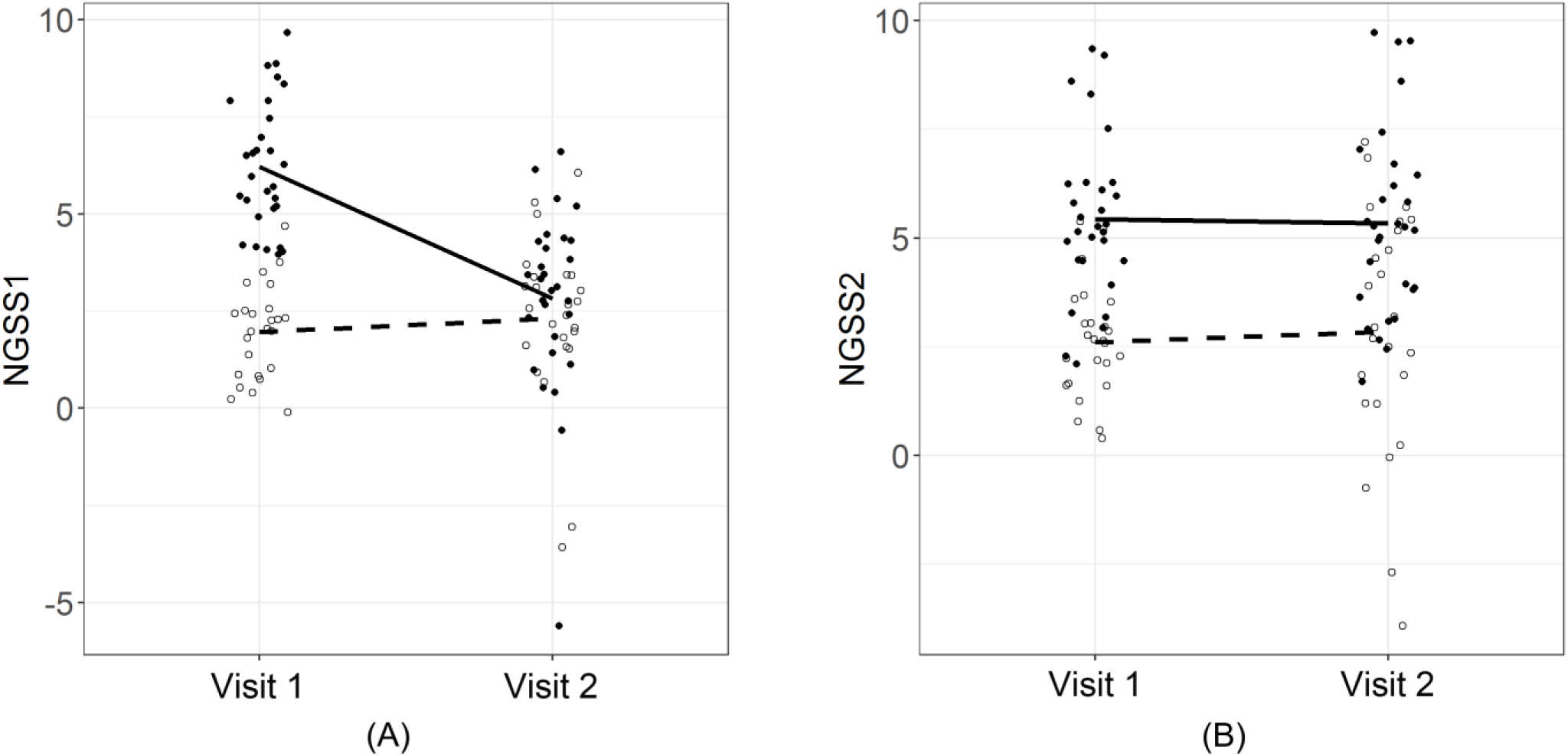
Paired comparisons between visit 1 and visit 2 using NGSS1 (panel (A)) and NGSS2 (panel (B)). A total of n=54 subjects with samples in both visits were used. Solid dots represent severe subjects and empty dots represent mild subjects. The solid line represents the mean trend of severe subjects and the broken line represents the mean trend for mild subjects. (A): At visit 1, there was a significant difference in mean NGSS1 between the severe (n=29) and mild (n=25) groups (6.22 vs. 1.96, p<0.001). Mean NGSS1 of the mild group was virtually unchanged between two visits (1.96 vs. 2.31, p=0.45). In comparison, mean NGSS1 of the severe group declined significantly at visit 2 (6.22 vs. 2.82, p<.001). (B): In contrast to NGSS1, the differences in NGSS2 was virtually unchanged between the two visits, due to the fact that NGSS2 were built with stable genes.

### Exploratory Association Analysis Based on Stable Nasal Genes

In the process of developing NGSS1 we observed that a large number of genes had expression levels that remained stable between the acute and convalescent visits. We speculated that a NGSS based on stable genes that were correlated with GRSS could potentially be predictive of disease severity prior to illness onset. Thus, we next developed an NGSS based on genes displaying stable expression across acute illness and convalescence in the 54 subjects with samples from both time points. Specifically, we included only genes whose mean expression levels correlated with disease severity during acute illness, and whose expression did not change significantly from the acute to convalescent stage.

We identified 2127 genes in subjects with mild illness and 1531 genes in subjects with severe illness, based on paired two sample t-test (p > 0.5) and fold change increases or decreases within 10%. Of the total 3658 genes, 689 stable genes were common in both groups (Figure 3A). A quality assurance analysis based on IQR showed that a small subset (n=14) of these genes had relatively small dynamic range in the combined dataset, and were excluded. We applied marginal screening based on Pearson correlation with GRSS to the remaining 675 stable genes and identified 44 marginally significant genes. As in developing NGSS1, we added 5 supplementary genes with strong marginal associations with GRSS. Model selection identified 13 genes as Model 4 (designated as **NGSS2**). The performance of NGSS2 is provided in Table 2 and illustrated in Figure 3B. NGSS2 showed a significant correlation with GRSS (ρ=0.741), and a CV accuracy of 84% (17 misclassifications out of 106 cases, Table 2). Of note, NGSS1 and NGSS2 do not contain any commonly selected gene, which is expected due to different screening criteria. Figure 2B shows that on average, NGSS2 did not change between visit 1 and visit 2, which is the key difference between these two classifiers. A full list of genes used in NGSS1 and NGSS2, as well as their estimated linear coefficients in the models, are listed in Supplementary Tables E2 and E3.

**Figure 3.**
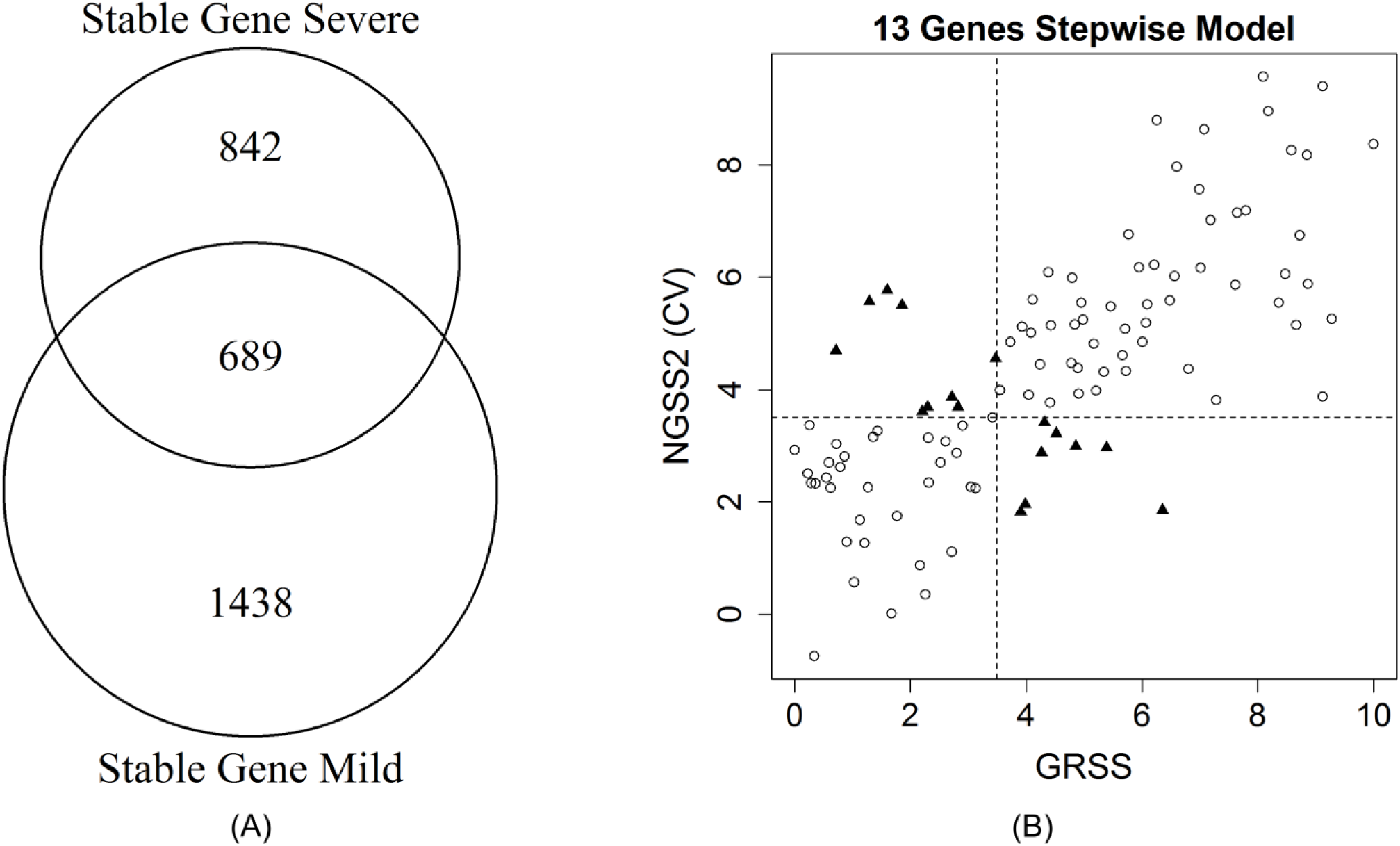
(A). Diagram indicating the stable genes for the mild (GRSS ≤3.5) and severe (GRSS >3.5) groups and the 689 intersecting stable genes common to both groups. (B). Correlating NGSS2 (severity score predicted by Model 4) with GRSS. Naïve Pearson correlation between GRSS and NGSS2 is ***ρ*** = **0.800**. Right: cross-validated Pearson correlation between GRSS and NGSS is ***ρ*** = **0.741**. Circles are subjects with correct cross-validated classification based on NGSS2; solid triangles are misclassified subjects.

## Discussion

Several approaches have been proposed for quantifying RSV disease severity in young infants.[4–13] A variety of clinical parameters have been included in several described severity scores, with incomplete agreement on the optimal factors to select.[14] One reason is that many clinical signs of RSV infection in young infants, including cutaneous oximetry, can fluctuate frequently and rapidly during the course of illness, making consistent assessment difficult. An objective biomarker reliably correlated with clinical severity could prove useful for clinical management and as a classifier and/or an outcome measure in vaccine or therapeutic trials.

Transcriptomic analysis of host cells has proven informative in the study of several respiratory viral infections, including RSV, with the emphasis on disease pathogenesis.[17–20] Unlike this report that focuses on nasal epithelial cell samples, most reports have described gene expression correlates of disease severity in peripheral blood mononuclear cells during infection since RSV pathogenesis is thought to be closely linked to the host’s immune response.[25] In two publications from the same group, RSV infection was associated with overexpression of innate immunity genes (neutrophil and interferon genes) and suppression of adaptive T and B cell genes. [17, 19] The investigators used the results to develop a gene-expression based illness score (designated Molecular Distance to Health [MDTH]) that was significantly correlated with a clinical disease severity score, duration of hospitalization and need for supplemental oxygen. Recently, Jong et al described an 84 gene signature that was highly predictive of RSV disease severity in infants.[16] Similarly, we reported that gene expression patterns in purified blood CD4 T cells during infection were correlated with clinical disease severity.[18] Gene expression results from nasal swabs collected from hospitalized infants during RSV infection have also been recently reported by another group, with differentially expressed genes correlated with clinical severity.[20]

In this report we describe the use of RNAseq analysis of gene expression data from nasal specimens collected during RSV infection to develop two nasal gene-expression severity scores (NGSS1 and NGSS2) that are highly correlated with a clinically derived disease severity score (GRSS). Although the nasal brush samples from the AsPIRES study were collected to investigate molecular pathways and disease mechanisms involved in pathogenesis (presented in a separate manuscript [26]), we also considered that the data could be useful for the development of a gene based biomarker of RSV severity. We used marginal screening of all genes followed by PCA analysis and step-wise model selection to develop NGSS1, a multivariate linear classifier of severity. In CV analysis, NGSS1 was strongly correlated with GRSS and was a relatively accurate classifier of binary disease severity. Furthermore, the score tracked well with clinical improvement 28 days after illness onset. Of particular note, we found that including uncorrelated supplementary genes enhances the accuracy of the models, and recommend this approach as a routine for future classification/prediction analyses based on high-throughput data with substantial correlation. As noted, in the population enrolled in our study the operating characteristics of NGSS1, including sensitivity, specificity, PPV and NPV, were quite good. However, it should be recognized that the proportion of mildly ill to severely ill subjects was determined by the recruiting strategy used, and that the PPV and NPV would vary depending on the population to which NGSS1 was applied.[21] If mildly ill subjects are increased by a factor of 3-5 this would reduce the PPV to 40-70% although the NPV would remain >90%.

Although the aim of this report is not to describe molecular mechanisms operative during RSV infection, it should be noted that the 41 NGSS1 genes include cytokines (TNFSF10, IL6, and CXCL2), extracellular matrix proteins (VIM, MMP19, RPS15A, FKBP1A, and VCAN), inflammation regulators (CXCL2, CD163), and components of various signaling processes (GNS, HAVCR2, PTPRC, CTSL, INHBA, IL6, MMP19, CXCL2, SLC39A8, CCDC80, VCAN, CD163). Some genes are only known to be involved in fundamental biological processes and are therefore novel in RSV research, including ST3GAL1 (a type II membrane protein) and ATP10B (ATPase Phospholipid Transporting 10B). Note that only two genes (TNFSF10, RABGAP1L) have been associated with disease severity in our recent study based on purified CD4 T cells.[18] In addition, IL-6 Signaling is the only significant canonical pathway identified from the CD4 T cells that contains an NGSS1 gene (IL6).

A unique and very preliminary result from our analysis is the development of NGSS2 using differentially expressed genes associated with GRSS that did not change between the acute and the convalescent time points. It is possible that these genes may simply be slow to return to baseline expression levels, in contrast to those genes selected for NGSS1. Although speculative, it occurred to us that “stable” genes might possibly be predictive of severity regardless of when a nasal sample was obtained, thus raising the possibility of infants at risk prior to or early in infection. While NGSS2 is slightly less accurate than NGSS1 in predicting GRSS during acute illness, the association between NGSS2 and GRSS is still relatively strong. Interestingly, the 13 NGSS2 genes were broadly related to cytoplasmic activities (EXOSC10, PLK2, PPIC, CLDN10, MAP3K13, MT1G, PXN), ATP binding (SEPHS2) and phosphoprotein regulation (BCKDK, PLK2, MAP3K13); activities that may be less directly responsive to acute RSV infection. These observations suggest that the best nasal transcriptome predictors of respiratory symptoms are not necessarily limited to those genes that directly regulate the immune response to RSV infection.

The use of nasal brush specimens for development of a severity biomarker in infants is attractive for a number of reasons. Nasal respiratory epithelial cells are the first cells infected and directly initiate early innate immune responses to RSV. The mucosa is also the site of migration of both innate and adaptive immune cells during infection. Importantly, we have shown that gene expression in nasal respiratory epithelial cells is highly concordant with published gene expression in lower respiratory tract epithelial cells, and thus should be a reasonable proxy for lung responses to RSV infection.[22] Of practical importance, collection of nasal epithelial cells is relatively non-invasive and simple to perform with minimal discomfort.

There are several important limitations to our study and conclusions. First, we do not have an independent cohort to validate our findings; the only publically available nasal gene expression data during RSV infection used microarray technology that did not identify many of the genes we identified by RNAseq. Due to the lack of independent samples for validation, we applied cross-validation techniques to prevent model overfitting and validate the accuracy of prediction for both NGSS1 and NGSS2 at the acute visit. CV estimator for prediction accuracy is known to be asymptotically unbiased [27] under very weak statistical assumptions, namely, the training and testing data are independent and identically distributed (which can even be relaxed further, see [28, 29]).

Additionally, we further validated the NGSS1 trained at the acute visit with the convalescence data, and the results conformed with our prediction remarkably well. Although the NGSS1 declined for the severely ill infants when clinical symptoms had resolved, it would be useful to determine if NGSS1 tracked closely over the full course of an illness. However, validation of our findings with an independent prospective cohort will be required. In addition, the results may not be valid for infants older than 10 months of age when infected with RSV, nor for infants with prematurity or other underlying medical conditions.

Another possible limitation is that all data used in these analyses were generated on the same technical platform and processed by the same team, therefore the validation results do not reflect the impact of “artifacts” in transcriptomic studies such as batch effects and platform differences, which can be reduced but not entirely eradicated by advanced normalization methods.[30–33]

Importantly, speculation that NGSS2 might predict disease severity prior to infection demands careful prospective validation. Finally, to extend the utility of time-intensive gene expression assays beyond a research tool and use it as a clinically useful biomarker of RSV disease severity, will require translation of these results to a rapid readily performed multiplex reverse transcription polymerase chain reaction (RT-PCR) assay, similar to those that have recently been developed for microbial diagnostics in respiratory secretions.[34]

In conclusion, we demonstrate that analysis of gene expression data obtained from an easily and safely obtained nasal brush specimen in young infants with acute RSV infection shows promise for development of composite molecular biomarkers that closely correlate with clinical severity score. Further studies to refine and validate the potential of predictive gene expression data from readily collected nasal samples are needed.

## Author Contributions

XQ, TJM, and EEW conceptualized the study. TJM, EEW, MTC, and CC designed the experiments. EEW, MTC, ARF and DJT developed the cohort, and collected the specimens and clinical data. LW, MNM, and XQ developed statistical models. JHW and AC facilitated data organization, management and analysis. LW, CC, MNM, CS, JH-W, AC, ARF, DJT, MTC, TJM, EEW, and XQ generated, analyzed and interpreted the data. LW, CC, MNM, CS, JH-W, AC, ARF, DJT, MTC, TJM, EEW, and XQ wrote and/or revised the manuscript. All authors read and approve the final manuscript.

## Acknowledgments

The transcriptional and microbiota data described in this manuscript are available in dbGaP (phs001201.v2.p1). An investigation of molecular pathways and disease mechanisms involved in pathogenesis is presented in Chu et al.[26]; a study of the chronology of airway microbiota dysbiosis associated with RSV infection, using the microbiota data set, is described in Grier et al.[35] This study is supported in part by Respiratory Pathogens Research Center (NIAID contract number HHSN272201200005C), and the University of Rochester CTSA award number UL1 TR002001 from the National Center for Advancing Translational Sciences of the National Institutes of Health. The content is solely the responsibility of the authors and does not necessarily represent the official views of the National Institutes of Health.

