## Supplementary Text for "Airway Gene-Expression Classifiers for Respiratory Syncytial Virus (RSV) Disease Severity in Infants"

### Supplementary Methods

#### Removing Batch-effects

Because the study spanned three winter seasons (October 2012 through April 2015) samples were processed in six library batches. This resulted in significant batch effects in total number of mapped reads (Supplementary Figure E1). In addition, analysis of variance (ANOVA) *F*-test with false discovery rate (FDR) controlled at 0.05 level, found 3,984 genes (28.8% of the reported transcriptome) had significantly different mean expressions across batches. Based on these observations, we applied ComBat to remove batch effects. After applying ComBat, none of the genes had significant batch effects based on ANOVA *F*-test, and pairwise correlation analysis showed that the average Pearson correlation between the original and ComBat processed data was 0.987. This suggests that ComBat only removed batch effects with minimum impact on the remaining information.

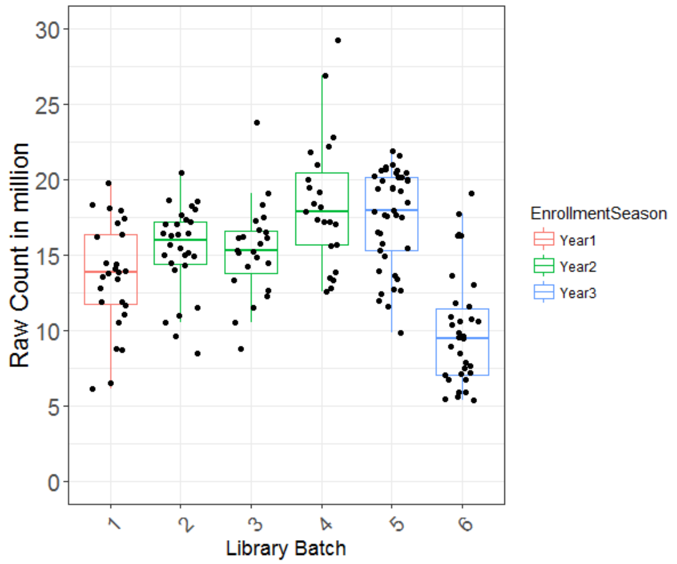


**Supplementary Figure E1**. Relationship between the total number of mapped reads, enrollment years, and batches in library preparation.

#### Model Developing and Cross-validation

***Identification of supplementary genes*.**

Both NGSS1 and NGSS2 were developed from top correlates (genes with significant marginal correlation with the GRSS) and some supplementary genes that contain information complementary to those top correlates. Specifically, we first performed principal component analyses (PCA) based on the decomposition of the correlation matrix of the top correlates (66 genes for NGSS1, and 44 genes for NGSS2), and used the leading principal component (PC1) to represent the collective information of those top correlates. Next, we conducted regression analyses with gene expressions as the response variables and the PC1 from the PCA as the covariate ($X_{ij} = \beta_{i0} + \mathrm{PC1} \cdot \beta_{i1} + \epsilon_{ij}$) for all genes except the 66 top correlates and recorded the residuals as $R_i = X_{ij} - \hat{\beta}_{i0} - PC1 \cdot \hat{\beta}_{i1}$. Here $X_{ij}$ was the expression of the $i$th gene sampled from the $j$th subject, and $R_i$ represented the information in $X_{ij}$ that was uncorrelated with PC1. Finally, we computed $\rho_i$, the Pearson correlation between $R_i$ and the GRSS, and select those genes with the largest $|\rho_i|$ as the supplementary genes (10 for NGSS1 and 5 for NGSS2).

***Model selection strategies.***

Two model selection strategies were used for developing predictive models based on multivariate regression analyses in which the GRSS was the response variable and genes were predictors. The first and primary strategy was a combination of stepwise model selection and an exhaustive search of a subset of all combinations of covariates. The second strategy is based on the elastic-net regularized regression (1, 2) and parameter refinement (3-5) based on the ordinary least-squares (OLS) criterion.

Strategy 1. We applied bi-directional stepwise model selection based on Akaike Information Criterion (AIC) to select an initial model. To further reduce model complexity, we selected a subset of *least informative* genes, defined as those with the smallest absolute values of the regression *t*-statistics in the initial model, and exhaustively searched the best sub-model of the initial model that do not include some of these 10 least informative genes (a total of 1023 combinations). Here the “best” sub-model was defined as the one with the smallest cross-validated residual sums of squares (CVRSS).

Strategy 2. As an alternative, we tried another model selection procedure based on regularized regression. Specifically, we first applied the elastic-net regularized regression, which uses both $L^1$ (LASSO) and $L^2$ (ridge) regression to produce a sparse regression model. The R package `glmnet` (2, 6) was used for this purpose. Regularization parameters were selected by an initial ten-fold cross-validation. After we obtain a sparse regression model, we re-estimate linear coefficients by OLS-based procedures to improve the accuracy of modeling fitting. The parameter refinement strategy can improve the prediction accuracy and was widely used in high-throughput data analysis (3-5).

Based on the empirical evidences in our study, we found that that Strategy 1 worked better than Strategy 2 for both NGSS1 and NGSS2, so we decided to use Strategy 1 for model selection and reported results based on Strategy 1 in the main text. A detailed comparison between these two strategies is provided in the following summary table.

**Supplementary Table E1**. Comparing the performance of two model selection strategies. For both NGSS1 and NGSS2, Strategy 1 worked better than Strategy 2 therefore it was selected in our study (see Table 2 in the main text).

| | number of genes selected | Naïve RSS | Naïve Correlation | Naïve misclassified subjects (out of 106) | CV RSS | CV Correlation | CV misclassified subjects (out of 106) |
|---|---|---|---|---|---|---|---|
| **NGSS1, Strategy1** | **41 genes** | **0.884** | **0.935** | **9** | **2.681** | **0.813** | **11** |
| NGSS1, Strategy2 | 20 genes | 1.725 | 0.869 | 17 | 2.617 | 0.800 | 23 |
| **NGSS2, Strategy1** | **13 genes** | **2.549** | **0.800** | **16** | **3.215** | **0.741** | **17** |
| NGSS2, Strategy2 | 20 genes | 2.606 | 0.795 | 16 | 3.816 | 0.688 | 20 |

In either case, the final predictor is a linear combination of gene expressions, which can be expressed as follows

$$NGSS_j = \hat{\beta}_0 + \sum_i^p X_{ij}\hat{\beta}_i,$$

where $\hat{\beta}_0$ stands for the intercept, $X_{ij}$ is the expression of the $i$th gene sampled from the $j$th subject, and $\hat{\beta}_i$ is the linear coefficient corresponding to the $i$th gene.

These estimates are summarized in Supplementary Tables E1 and E2.

<u>*Cross-validation*</u>. For both strategies, we evaluate the performance of the fitted models by leave-one-out cross-validations. The residual sum of squares (RSS), Pearson correlation between the actual and predicted GRSS, and prediction accuracy based on the 3.5 cutoff for mild versus severe symptoms, are recorded in these CV studies.

**Supplementary Table 2**. List of genes used in NGSS1 (Model 2) and the corresponding linear coefficients. Source: "Sig." one of the original 66 significant genes; "Supp.": a supplemental gene.

| Gene symbol | Estimated linear coefficient ( $\hat{\beta}$ ) | Marginal p-value | Source |
|---|---|---|---|
| Intercept ($\hat{\beta}_0$) | 0.703269912 | 6.85E-01 | - |
| ST3GAL1 | 0.059521015 | 1.21E-03 | Sig. |
| VIM | 0.003782824 | 1.41E-01 | Sig. |
| VCAN | -0.010671636 | 3.60E-02 | Sig. |
| CXCL2 | 0.012153063 | 2.36E-02 | Sig. |
| PTPRC | -0.009655874 | 1.31E-02 | Supp. |
| FKBP1A | -0.043599137 | 1.05E-05 | Sig. |
| pk | 0.122119918 | 8.70E-06 | Sig. |
| CCDC80 | -0.007741724 | 2.38E-02 | Sig. |
| HMOX1 | -0.02629094 | 8.67E-02 | Sig. |
| NKG7 | -0.001696102 | 5.20E-01 | Sig. |
| LPXN | 0.018386158 | 2.95E-01 | Sig. |
| PHACTR2 | 0.069346751 | 7.53E-03 | Sig. |
| TIA1 | -0.033380848 | 2.83E-02 | Sig. |
| ATP10B | 0.027307907 | 1.97E-01 | Sig. |
| TNFSF10 | -0.0018723 | 2.96E-02 | Sig. |
| INHBA | -0.022899548 | 3.61E-02 | Sig. |
| MMP19 | 0.021311619 | 1.42E-01 | Sig. |
| SMUG1 | -0.012079396 | 2.43E-01 | Supp. |
| MPP1 | -0.010764038 | 2.10E-01 | Sig. |
| RPS15A | 0.004696851 | 8.16E-05 | Supp. |
| CTSL | -0.007780092 | 7.84E-05 | Sig. |
| HAVCR2 | 0.073470635 | 6.43E-06 | Sig. |
| GNS | 0.105879463 | 1.13E-04 | Sig. |
| IL6 | 0.012406829 | 2.53E-02 | Sig. |
| SLC39A8 | -0.034782958 | 8.17E-02 | Sig. |
| PTPN7 | -0.036920209 | 1.85E-02 | Sig. |
| RABGAP1L | -0.007262571 | 2.92E-01 | Sig. |
| SLC7A7 | 0.02358896 | 1.27E-01 | Sig. |
| ITGA5 | -0.033753825 | 1.75E-02 | Sig. |
| PPM1M | 0.053787695 | 9.91E-03 | Sig. |
| ARFIP1 | 0.03076839 | 1.21E-02 | Sig. |
| MFSD4 | -0.007710719 | 1.34E-01 | Sig. |
| CD163 | 0.007982385 | 6.33E-03 | Sig. |
| SHMT2 | 0.017423769 | 1.61E-01 | Supp. |
| SERHL2 | 0.04069629 | 3.42E-05 | Supp. |
| MAFB | 0.080380399 | 1.53E-02 | Sig. |

| | | | |
|---|---|---|---|
| C10orf128 | 0.037369718 | 5.12E-02 | Sig. |
| FTLP3 | 0.012010208 | 5.11E-02 | Sig. |
| RP11-206L10.8 | -0.048018562 | 8.85E-04 | Sig. |
| N4BP2L2 | -0.011088274 | 1.28E-02 | Sig. |
| VCAN-AS1 | -0.014399275 | 1.31E-01 | Sig. |

**Supplementary Table E3**. List of genes used in NGSS2 (Model 4) and the corresponding linear coefficients. Source: "Sig.": one of the 44 original significant genes; "Supp.": a supplemental gene.

| Gene symbol | Estimated linear coefficient ( $\hat{\beta}$ ) | Marginal p-value | Source |
|---|---|---|---|
| Intercept ($\hat{\beta}_0$) | 5.47173684 | 9.43E-05 | - |
| EXOSC10 | -0.05176355 | 7.20E-03 | Sig. |
| PPIC | -0.05462252 | 1.94E-05 | Sig. |
| CCNI | 0.01055988 | 7.07E-06 | Sig. |
| BCKDK | 0.06716161 | 5.04E-04 | Sig. |
| MAP3K13 | -0.01209091 | 1.26E-02 | Sig. |
| MT1G | 0.00242952 | 5.62E-02 | Sig. |
| APOC1 | -0.00404016 | 4.64E-02 | Sig. |
| QTRTD1 | -0.04002261 | 7.18E-03 | Sig. |
| DDRGK1 | -0.02822089 | 2.94E-02 | Sig. |
| SEPHS2 | 0.07799082 | 2.65E-03 | Sig. |
| PLK2 | 0.03691206 | 2.02E-02 | Sig. |
| CLDN10 | 0.01013402 | 4.33E-04 | Supp. |
| PXN | -0.07814369 | 1.29E-04 | Supp. |
